# Chronic nicotine treatment enhances cognition and reduces neuroinflammation in the gp120 transgenic mouse model of neuroHIV

**DOI:** 10.1101/2025.05.03.651604

**Authors:** Samantha M. Ayoub, Tyler Dexter, Michael Noback, Melissa Flesher, Cris Achim, Jerel Adam Fields, Arpi Minassian, Arthur L. Brody, Jared W. Young

**Affiliations:** Department of Psychiatry, University of California San Diego, La Jolla, CA; Research Service, VA San Diego Healthcare System, San Diego, CA

**Keywords:** Smoking, risk-taking, reinforcement learning, Iba-1, motivation, cross-species task

## Abstract

**Rationale:** Antiretroviral development has improved the longevity of people with HIV (PWH), but many experience impaired cognition potentially due to neuroinflammation. PWH smoke cigarettes at higher rates than the general population, possibly for self-medication given cognitive-enhancing and anti-inflammatory properties of nicotine, the primary psychoactive ingredient cigarette smoke. We hypothesized that chronic nicotine would improve cognition in a mouse model of HIV, gp120 transgenic (Tg) mice, and reduce neuroinflammation.

**Methods:** Male and female gp120-Tg mice (n=64) and littermate controls (n=67) were operantly trained then tested for effortful motivation in the progressive ratio breakpoint task (PRBT). Mice were counter-balanced into three groups for saline or nicotine minipump implantation (0, 14 or 40 mg/kg/day) then retested 25 days later in the PRBT, probabilistic reversal learning task (PRLT – reinforcement learning and cognitive flexibility), and Iowa Gambling Task (IGT – risk-based decision-making), with a subset tested for neuroinflammation (Iba-1 levels).

**Results:** Gp120-Tg mice exhibited worse PRLT performance, attenuated by nicotine. Furthermore, nicotine selectively optimized their response strategies in the PRLT and IGT, increasing loss sensitivity, shifting animals towards “safer” responses. No motivation effects were observed. Nicotine also reduced Iba-1 expression, suggesting that its cognitive-enhancing effects may relate to reduced neuroinflammation.

**Conclusion:** Gp120-Tg mice exhibited deficits in the PRLT, which are attenuated by chronic nicotine. Furthermore, nicotine improved reinforcement learning and risky decision-making supporting its therapeutic potential for cognitive deficits in PWH, possibly via reducing neuroinflammation. With potential negative consequences of long-term nicotine use, future studies should determine its mechanism of action to develop more targeted therapeutics.

## Introduction

Nearly half of people living with HIV (PWH) exhibit some form of neurocognitive impairment (NCI) [1, 2], for which no targeted treatments exist. Current antiretroviral therapies (ART) minimize the likelihood of HIV development to Acquired Immunodeficiency Syndrome (AIDS), but NCI is still observed in virally-suppressed PWH [3]. Thus, PWH now live longer and generally healthier lives, with the complication of persisting NCI.

When compared to the general population, PWH smoke nicotine-based products at a higher rate and are also less likely to quit [4, 5]. The impact of this comorbid substance use on HIV-associated NCI is not well understood, however. On one hand, smoking puts PWH at a higher risk of smoking-related illnesses and may reduce ART effectiveness [6]. On the other hand, numerous experimental human and animal studies suggest that nicotine, the primary psychoactive ingredient in e-vapes and cigarettes, has cognitive enhancing properties [7–10]. Nicotine also has anti-inflammatory efficacy [11]. Since neuroinflammation is observed in PWH and proinflammatory markers are linked to NCI in PWH [12–16], nicotine may benefit HIV-associated NCI.

Some clinical evidence suggests smoking negatively impacts cognition in PWH. For example, current smoking status shared an inverse relationship with global cognition, as well as learning, and memory performance in PWH [17]; however, this effect disappeared when accounting for education and the presence of comorbid infections (Hepatitis C). Another group reported working-memory and processing-speed deficits in a group of HIV+ smokers, relative to HIV-smokers, though the exclusion of non-smoking controls limits interpretation of these findings as it could simply reflect HIV-induced NCI [18]. Nonetheless, two well-controlled studies reported interactions between HIV and smoking status which revealed poorer global cognition and decision-making function in PWH who smoke, relative to non-smokers and healthy controls [19, 20]. Importantly however, these latter studies did not report the time course of testing vs. last smoking exposure. Such information is vital because nicotine withdrawal-induced cognitive deficits cannot be ruled out from causing the observed adverse effects [21–26]. Timing since last use may explain why past, but not current, smoking status was associated with better cognition in PWH [27], potentially resulting from withdrawal-induced cognitive deficits relevant to current smokers. In a recent study from our group where participant smokers were allowed to use just prior to cognitive testing, PWH who smoke exhibited enhanced cognitive control and reduced positron emission tomography (PET)-measured neuroinflammation relative to non-smoking PWH and healthy control participants [15]. Others have reported no effect of cigarette smoking on cognition in PWH [28]. Hence, clinical data remains equivocal as to the effect of smoking on HIV-associated NCI, with interpretations complicated by smoking initiation, recent usage, and dosing levels, not always reported.

Delineating the effect of smoking on cognition in PWH may be better answered experimentally using animal models of neuroHIV to allay the potential confounds. The importance of such factors is highlighted by the Bryant et al. (2013; [17]) findings that the adverse effect of smoking on cognition in PWH was negated once data were corrected for educational and comorbidity differences across groups. Such variables are easier to control for in animal models, which also enable identical drug exposures (length of use, dosage) across groups, another factor inherently difficult to measure and/or control for in human populations. Further, the causality of human findings remains unclear given their associative nature, i.e., do differences in cognitive function precede smoking initiation? Animal testing enables experimentally defining more causal mechanisms informing directionality of effects. Finally, while nicotine is thought to be anti-inflammatory and support cognition, cigarettes contain additional harmful ingredients that may independently impact cognition and inflammatory processes [as reviewed by 29], underscoring the importance of parsing apart the individual effects of nicotine use in this population.

Well-controlled studies increase the ability to delineate the potential effects of nicotine on HIV-associated NCI. In general, interactions between HIV-relevant manipulations and nicotine effects have been observed on cognitive and behavioral outcomes, though data are limited. For example, chronic nicotine reduced contextual fear memory deficits in a model of HIV that express 7 of 9 HIV proteins, the HIV-1 transgenic (Tg) rats, and reversed HIV-related alterations in synaptic plasticity gene expression in the brain, while impairing such memory in control rats [30]. In contrast, chronic nicotine had modest procognitive effects in control rats without affecting HIV-1Tg rats in spatial learning in a water maze [31]. Taken together, the impact of nicotine on cognitive function in HIV may be task-dependent, consistent with cannabis effects in PWH [32]. The effects of nicotine in neuroHIV animal models however, are relatively unexplored in cross-species cognitive tasks and require further assessment.

HIV glycoprotein (gp)120 Tg mice are one of the first animal models of neuroHIV which overexpress the viral protein gp120 responsible for HIV penetration into cells [33]. Gp120-Tg mice effectively model behavioral/cognitive abnormalities [34–36] and neuropathological features [33, 35, 37, 38] including PET-measured neuroinflammation [39], akin to PWH, suggesting the gp120 protein as an important contributor to HIV-associated symptomatology, including NCI. Importantly, alpha7 nicotinic acetylcholine receptors (α7 nAChR), to which nicotine binds, are upregulated in the striatum of gp120-Tg mice [40, 41] and on immune cells in post mortem brains of PWH [42], suggesting this model is particularly relevant for studying the interactions of HIV and nicotine in cognition. Consistently, acute gp120 increased α7 nAChRs expression *in vitro* [40]. To our knowledge, only one study has explored the cognitive impact of nicotine in gp120-Tg mice, however. In this study, subchronic nicotine injections attenuated gp120-induced motor coordination dysfunction, as well as abnormal event related potentials in the gp120-Tg mouse model [43], suggestive of procognitive effects. Since HIV-associated NCI is proposed to arise from neuroinflammation, while proinflammatory markers in the CNS of PWH are linked to NCI [12–14, 16], nicotine may reduce neuroinflammation in gp120-Tg mice and enhance cognition.

To address the current literature gap, the interactive and independent effects of chronic nicotine exposure and HIV-relevancy on cognition and brain microglial activation were tested in the gp120-Tg mouse model of HIV. Importantly, cognitive processes were tested using cross-species tasks to enhance translatable findings [44–46] including the progressive ratio breakpoint task (PRBT; to measure effortful motivation), the probabilistic reversal learning task (PRLT; to measure learning and cognitive flexibility), and the Iowa Gambling Task (IGT; to measure risk-based decision-making). The brains of some mice were collected to assess altered microglia activation (Iba-1) as a proxy for neuroinflammation. We hypothesize that that chronic nicotine treatment will improve the cognitive performance of gp120-Tg mice in these tasks via reducing neuroinflammation.

## Materials and Methods

### Animals

Adult gp120-Tg (n=64) and wildtype (WT; n=66) littermate control mice were used for the current studies (N=130; ♀=69, ♂=61; see Table 1). Animals were aged 12 weeks and weighed 19-28 g at the beginning of the study. Mice were generated from a mixed C57BL/6 x Sv129 (SJL/BL6/129) background [47, 48] with gp120 expression occurring in astrocytes under the control of a modified murine glial fibrillary acidic protein (GFAP) promoter [33]. For all experiments, Tg mice on a mixed BL6/129 background were crossed with BDF1 mice (both male and female) obtained from Charles River, and Non-Tg WT littermates were used as controls. Genotypes were confirmed via PCR analysis of tail DNA (Transnetyx). During training and testing mice were food restricted to 85% of their free-feeding weight, with access to water provided *ad libitum*. Mice were housed in groups of 1–4 per cage in a climate-controlled animal colony on a reversed day/night cycle (lights on/off at 7:00/19:00), and testing conducted between 12:00 and 6:00 PM. Mice were maintained in a University of California San Diego (UCSD) animal facility which meets all federal and state requirements for animal care, and all procedures were approved by the Institutional Animal Care and Use Committee at UCSD.

**Table 1.** Sample sizes by sex, genotype, and drug assignment.

| **Sex** | **Genotype** | **Nicotine** | **Sample size** |
| --- | --- | --- | --- |
| **Female**<br>♀ | **WT littermates** | Vehicle | 14 |
|  |  | 14 mg/kg/day | 10 |
|  |  | 40 mg/kg/day | 9 |
|  | **Gp120-Tg** | Vehicle | 13 |
|  |  | 14 mg/kg/day | 11 |
|  |  | 40 mg/kg/day | 12 |
| **Male**<br>♂ | **WT littermates** | Vehicle | 10 |
|  |  | 14 mg/kg/day | 11 |
|  |  | 40 mg/kg/day | 12 |
|  | **Gp120-Tg** | Vehicle | 9 |
|  |  | 14 mg/kg/day | 11 |
|  |  | 40 mg/kg/day | 8 |

### Apparatus: 5-hole operant chamber

Sound-insulated 5-hole operant chambers (25×25×25 cm; Med Associates, Inc., St. Albans, VT) were utilized. Each chamber had a house light and fan, with an array of 5 square holes (2.5×2.5×2.5 cm, 2.5 cm above the grid floor) arranged horizontally on a curved wall opposite the liquid delivery magazine (Lafayette Instruments, Lafayette, IN). Each hole had a light-emitting diode (LED) at the back and infrared beams, mounted vertically 3 mm from the opening to detect responses. The food delivery magazine contained a well for liquid reinforcement (strawberry Nesquik® plus non-fat milk, 30 μL), delivered by a peristaltic pump (Lafayette Instruments, Lafayette, IN), with an LED at the top. The magazine also contained an infrared beam to detect head entries. The control of stimuli and recording of responses were managed by a SmartCtrl Package 8-In/16-Out with additional interfacing by MED-PC for Windows (Med Associates, Inc.) using custom programming.

### Training and Testing

Training consisted of two phases, consistent with previous reports [49]. Mice were first conditioned to associate an illuminated magazine with reward delivery during a 20 min session wherein ∼30 µl strawberry milkshake was dispensed on a 15 sec fixed interval schedule. Mice were maintained on this program until at least 60 reward collections were made across two consecutive days. Next, animals were required to make an operant response (nosepoke) into one of five illuminated choice apertures to earn reward on a fixed-ratio 1 (FR-1) schedule of reinforcement during daily 30 min sessions (Figure 1A). Importantly, after 5 continuous responses in a single aperture the aperture was disabled to prevent the development of side and/or aperture biases. Disabled apertures were reactivated following 2 responses in other apertures. Criterion for completing this training phase was set at 70 correct nose pokes per session for two consecutive sessions, with stable FR-1 responding over 4 consecutive days, prior to baseline assessment in the PRBT. Each testing day was separated by a FR-1 schedule of reinforcement training day which utilized the number of choice apertures required for the respective task (PRBT n=1; PRLT n=2; IGT n=4). This was done to familiarize mice with the apertures to be used in the subsequent test, and to re-stabilize FR-1 activity between tests.

**Figure 1.**
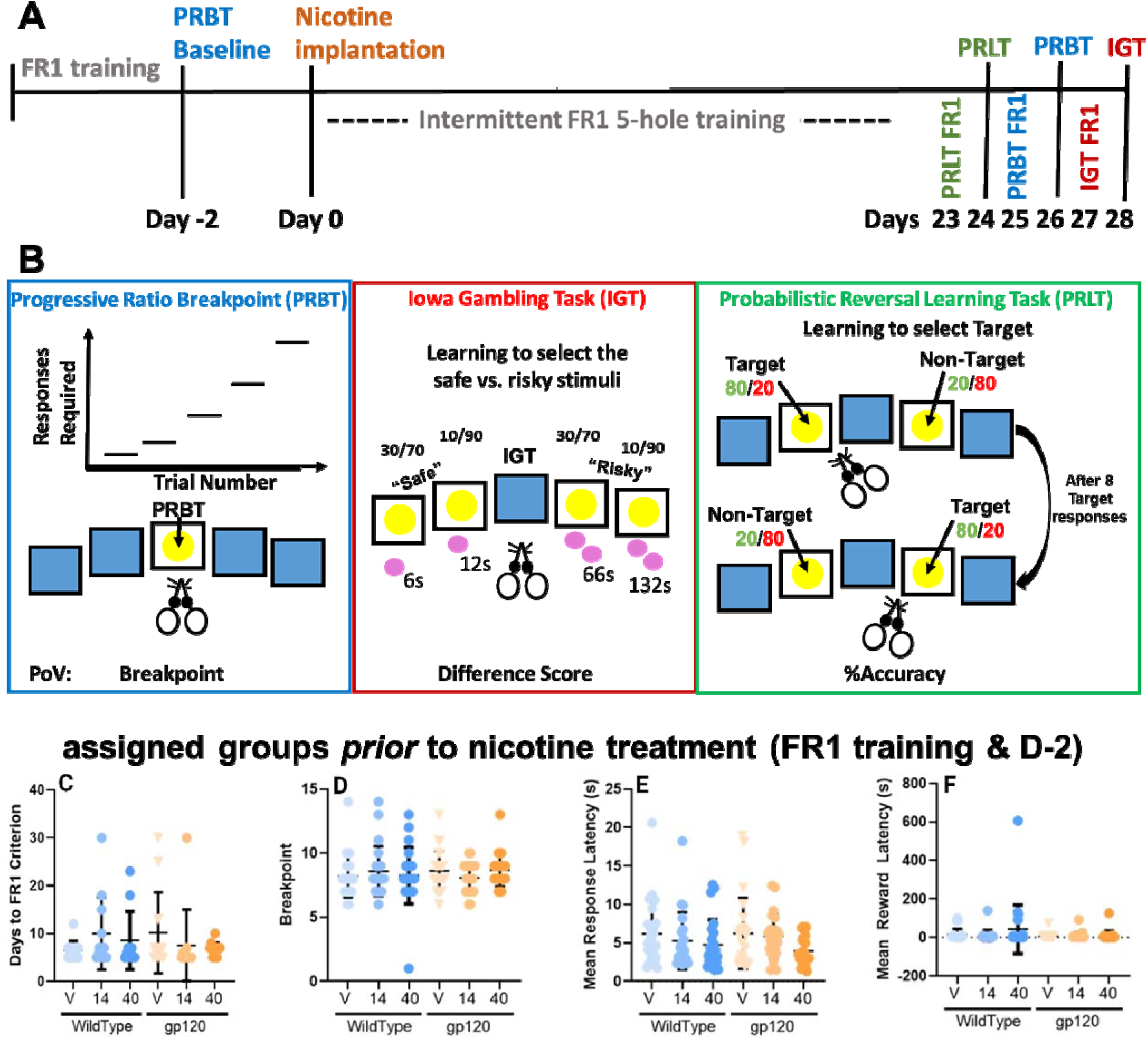
Timeline and schematics, with equal performance of learning and motivation in gp120-Tg mice. **A.** Experimental timeline showing initial Fixed-ratio 1 (FR-1) training, responding in one of 5 single lit holes used throughout testing for 1 reward prior to baseline assessment in the PRBT (Day −2), nicotine implantation (day 0), and in FR-1 for specific holes prior to testing (Days 23-28). **B.** Schematic and primary outcome variables of the 3 tasks utilized with PRBT requiring increasing response rates over time for the same 1 reward (left), IGT choosing between safe and risky options (middle), and PRLT, selecting the option more likely to result in a reward (target, 80/20), than a punishment (non-target; 20/80). **C-F.** No effects of genotype, or assigned dose, nor their interaction were observed for the days to reach criterion **(C)**, breakpoint **(D)**, mean response latency **(E)**, or mean reward collection latency **(F)**. Data presented as individual data-points, plus mean ± standard error of the mean (S.E.M).

### Progressive Ratio Breakpoint Task (PRBT)

The PRBT has long been used to assess effortful motivation for rewards, whether drug or food, which requires animals to consistently nosepoke in the central hole (Figure 1B). The number of hole pokes required to gain a reward increased according to the following exponential progression: 1, 2, 3, 6, 9, 12, 15, 20, 25, 32, 40, 50…, derived from the formula (5 x ^e0.2n^) – 5) rounded to the nearest integer where n is the position in the sequence of ratios. Each trial was repeated for 3 trials to ensure continuity of nosepoking as described previously [50, 51]. The session continued for 60 min or until 5 min had passed without a single nose poke. The main outcome variable was the breakpoint, defined as the last ratio to be completed before the session end. Mean response latency and mean reward latency were also calculated.

### Probabilistic Reversal Learning Task (PRLT)

The PRLT is a 60 min within-session reversal learning paradigm that measures both initial and reversal learning about probabilistic reinforcement (Figure 1B). This task utilized 2 of the 5 choice apertures, evenly spaced apart. Trials were initiated following a nosepoke into the reward magazine which triggered the illumination of two choice apertures following a 5 sec ITI. The target hole provided a high probability of reward (80%) and low probability (20%) of ‘punishment’ (4 sec house light illumination). Although not strictly a punishment, the illuminated house light is used to indicate the lack of reward for the selection made – similar to human PRLTs wherein the subject is simply informed they were ‘wrong’ via feedback [(e.g., red frowny face; 52, 53]. Instead, the nontarget hole provided a low probability of reward (20%) and high probability of punishment (80%). After 8 consecutive target hole responses, the reward and punishment contingencies at the target and nontarget locations were switched. All outcomes variables of the task are described in Table 2. The primary outcome measures included the number of switches achieved and the percentage (%) of Target responses. The secondary outcome measures included %premature responses, total trials, win-stay and lose-shift behaviors at both target and nontarget locations, choice latencies at both target and nontarget locations, reward latency, time-out responses, and preservative responses.

**Table 2.** Main Outcome Variables from the PRLT.

|  |  |  |
| --- | --- | --- |
| **Primary Outcomes** | Switches | Number of reversals achieved after 8 consecutive responses in the Target location |
|  | %Target Responses | Percentage of trials with Target location selection |
| **Secondary Outcomes** | %premature Responses | Percentage of trials with a choice response made prior to target/non-target stimuli presentation |
|  | Total Trials | Choice and premature trials summed |
|  | Target Win-Stay Ratio | Reward sensitivity. Tendency to reselect the Target location after receiving a reward from making a Target location selection in the previous trial |
|  | Target Lose-Shift Ratio | Punishment sensitivity. Tendency to shift choice selection to the NonTarget location after receiving punishment from making a Target location selection in the previous trial. |
|  | NonTarget Win-Stay | Reward sensitivity. Tendency to reselect the NonTarget location after receiving a reward from making a NonTarget location selection in the previous trial. |
|  | NonTarget Lose-Shift Ratio | Punishment sensitivity. Tendency to shift choice selection to the Target location after receiving punishment from making a Target location selection in the previous trial. |
|  | Mean Target Latency | Latency to select (nosepoke) the Target location |
|  | Mean NonTarget Latency | Latency to select (nosepoke) the NonTarget location |
|  | Mean Premature Latency | Latency to make a premature response |
|  | Mean Reward Latency | Latency to collect reward |
|  | Time-out Responses | All responses made during time-out punishment |
|  | Reward Preservation | Repetitive responses made into any choice aperture following the delivery of reward |
|  | Punish Preservation | Repetitive responses made into any choice aperture following the delivery of punishment |

### Iowa Gambling Task (IGT)

The IGT was initially designed to assess real-world decision-making under uncertain risk for humans and translated for animal use in a single session [as previously described; 50, 54]. The test lasted up until 400 trials were completed or 60 minutes elapsed. During the IGT, mice were presented with 4 options (Figure 1B). Selecting one of these options resulted in either a reward or punishment at the level and probabilities provided in Figure 1B. Specifically, selecting the two left holes (“risky”) could result in 2 rewards (∼50 µl), or long time-out punishments (66 or 132 s) at a probability of 50/50 or 25/75 respectively. Selecting the two right holes (“safe”) could result in 1 reward (∼25 µl), or short time-out punishments (12 or 6 s) at a probability of 75/25 or 50/50, respectively. The risky and safe holes were on the left or right kept consistent per mouse throughout, with the safe side opposite the side-bias displayed by the mice during training. The main outcome was the degree of safe vs. risk-preference, quantified using a difference score (no. of safe – risky responses). Secondary outcomes included rewards earned, punishment duration experienced, %omission trials, %premature trials, and latencies for choice behavior and reward collections.

### Drug

(−)Nicotine hydrogen tartrate (Sigma-Aldrich, St. Louis, MO) was dissolved in sterile 0.9% saline solution and pH adjusted to 7±0.5 with sodium hydroxide (Sigma-Aldrich). Nicotine was infused through subcutaneous osmotic minipumps at concentrations of 14 and 40 mg/kg/day (Model 2004, ALZET, Palo Alto, CA). Doses were chosen based on previous reports of nicotine’s behavioral and neurobiological effects in mice [55, 56].

### Osmotic mini pump implantation surgery

The ALZET mini-osmotic pump Model 2004 has a reservoir volume of 200 µl and delivers solutions at a pumping rate of 0.25 µl/h (±0.05 µl/h). Pumps were filled and primed in 0.9% saline solution at room temperature for 40-48 h before insertion. Mice were anesthetized with isoflurane (1-3% in oxygen). Before inserting the minipump, the area around the back of the neck was shaved and sterilized with betadine. An incision was made and a pouch large enough for the pump was blunt dissected into the back using scissors. The pre-filled pump was inserted into the pouch with the flow modulator directed posteriorly, away from the wound. The incision was closed with 9 mm wound clips (MikRon Precision, Inc., Gardena, CA) and baytril (5 mg/kg) and flunixamine (2.5 mg/kg) were injected subcutaneously to minimize chances of infection and alleviate pain, respectively.

### Immunohistochemistry of brain sections

Briefly, as previously described [57], mouse hemibrain tissue sections were deparaffinized using xylene followed by rehydration in serial ethanol and water solutions. Next, tissue sections were treated for 30 min with 3% hydrogen peroxide/phosphate-buffered saline (PBS) and then incubated for 30 min with 2.5% normal serum, corresponding to the host species for the secondary antibody. Tissue sections were then incubated with anti-IBA1 (Wako, cat. no. 019–19741; 1:100 in PBS) for 2 h at room temperature in a hydration box. Subsequently, tissue sections were washed with 0.1% Tween 20/PBS, before 30 min incubation with horse anti-rabbit IgG peroxidase-polymer secondary antibody (ImmPRESS, Vector Laboratories, Burlingame, CA, USA). After the tissues were washed with 0.1% Tween 20/PBS, the signals were developed with diaminobenzidine (ImmPACT DAB peroxidase substrate, Vector Laboratories) for 5 min. The immunostained sections were then dehydrated via serial ethanol and water solutions, de-waxed with xylene, and mounted using Cytoseal 60 (ThermoScientific). For the negative control, the primary antibody was omitted.

Subsequently, immunostained sections were scanned using a microscope slide scanner (Aperio ScanScope GL, Leica Biosystems, Buffalo Grove, IL, USA) equipped with a 20 × objective lens (yielding the resolution of 0.5 μm per pixel). Assessment of levels of IBA1 immunoreactivity was performed using the Aperio ImageScope software. For each case a total of three sections (5 images per section) were analyzed to estimate the average optical density of immunolabelled cells per unit area (mm^2^). Corrected optical density was calculated by subtracting the background optical density of the negative control (obtained from tissue sections immunostained in the absence of primary antibody) from the optical density of the immunostained sections.

### Statistical analyses

The primary outcomes from each task and histology were analyzed using analyses of variance (ANOVA), with genotype and sex as between-subjects factors during baseline, with nicotine as an additional factor. When data were analyzed across time, time was included as a within-subjects factor). Alpha was set at p<0.05, though trend effects (p<0.1) were also reported when observed. Secondary outcome variables were analyzed similarly but with Bonferroni corrections included for multiple assessments. Tukey *post hoc* analyses were conducted on all significant effects and interactions. The impact of nicotine on gp120-Tg mice alone were also analyzed on primary outcome variables based on our *a priori* hypotheses. One nicotine-treated WT mouse did not record any data during PRBT testing and was removed from the analysis. All statistical analyses were conducted using SPSS v 27 (Chicago, IL).

## Results

### Pre-nicotine exposure baseline performance

All mice were assigned to nicotine conditions prior to mini-pump implantation. Baseline performance on the FR-1 and PRBT tests was compared across genotypes, and future drug condition (“assigned dose”) was also included as a factor to ensure there were no off-target differences between groups at baseline (Figure 1 A-B). For initial FR-1 training, we did not observe effects of genotype or assigned nicotine dose (Fs<0.804, ns; Figure 1C). A main effect of sex was observed [F(1,118)=11.188,*p*=0.001], revealing that males attained FR-1 criterion faster than females [6.11 (±0.585) vs. 8.57 (±0.567) respectively]. No sex*gene, sex*assigned dose, or sex*gene*assigned dose, interactions were observed (Fs<2.626, ns). Mice were then tested on the PRBT.

<u>Breakpoint:</u> No main effect of genotype, assigned dose, nor interactions between sex*genotype, sex*assigned dose, genotype*assigned dose, or sex*genotype*assigned dose were observed (Fs<0.45, ns; Figure 1D). We did observe a trend effect of sex [F(1,118)=3.90,*p*=0.051], revealing that male mice exhibited a higher breakpoint than female mice (Supplementary Figure 1A).

<u>Mean Response Time:</u> A significant effect of sex was observed [F(1,118)=9.60,*p*=0.002], revealing that male mice exhibited faster response times than female mice (Supplementary Figure 1B). No main effect of genotype, assigned dose, nor interactions between sex*genotype, sex*assigned dose, genotype*assigned dose, or sex*genotype*assigned dose were observed (Fs<2.3, ns; Figure 1E).

<u>Mean Reward Latency:</u> A significant effect of sex was observed [F(1,118)=4.40,*p*=0.038], revealing that male mice exhibited slower latencies to collect their rewards than female mice (Supplementary Figure 1C). No main effect of genotype, assigned dose, nor interactions between sex*genotype, sex*assigned dose, genotype*assigned dose, or sex*genotype*assigned dose were observed (Fs<1.35, ns; Figure 1F).

### Effects of chronic nicotine exposure

#### Progressive Ratio Breakpoint Task

Mice were first tested on the PRBT (Figure 2). <u>Breakpoint:</u> A significant effect of sex was observed [F(1,117)=7.15,*p*=0.009], revealing that male mice exhibited a higher breakpoint than female mice (Supplementary Figure 1D), as seen at baseline testing. No main effect of genotype, assigned dose, nor interactions between sex*genotype, sex*assigned dose, genotype*assigned dose, or sex*genotype*assigned dose were observed (Fs<1.08, ns; Figure 2A). <u>Mean Response Time:</u> A significant effect of sex was observed [F(1,117)=8.87,*p*=0.004], revealing that male mice exhibited faster response times than female mice (Supplementary Figure 1E), as seen at baseline testing. No main effect of genotype, assigned dose, nor interactions between sex*genotype, sex*assigned dose, genotype*assigned dose, or sex*genotype*assigned dose were observed (Fs<0.55, ns; Figure 2B). <u>Mean Reward Latency:</u> A significant effect of sex was observed [F(1,117)=18.95,*p*<0.001], revealing that male mice collected their rewards slower than female mice (Supplementary Figure 1F), as seen at baseline testing. No main effect of genotype, assigned dose, nor interactions between sex*genotype, sex*assigned dose, genotype*assigned dose, or sex*genotype*assigned dose were observed (Fs<1.80, ns; Figure 2C).

**Figure 2.**
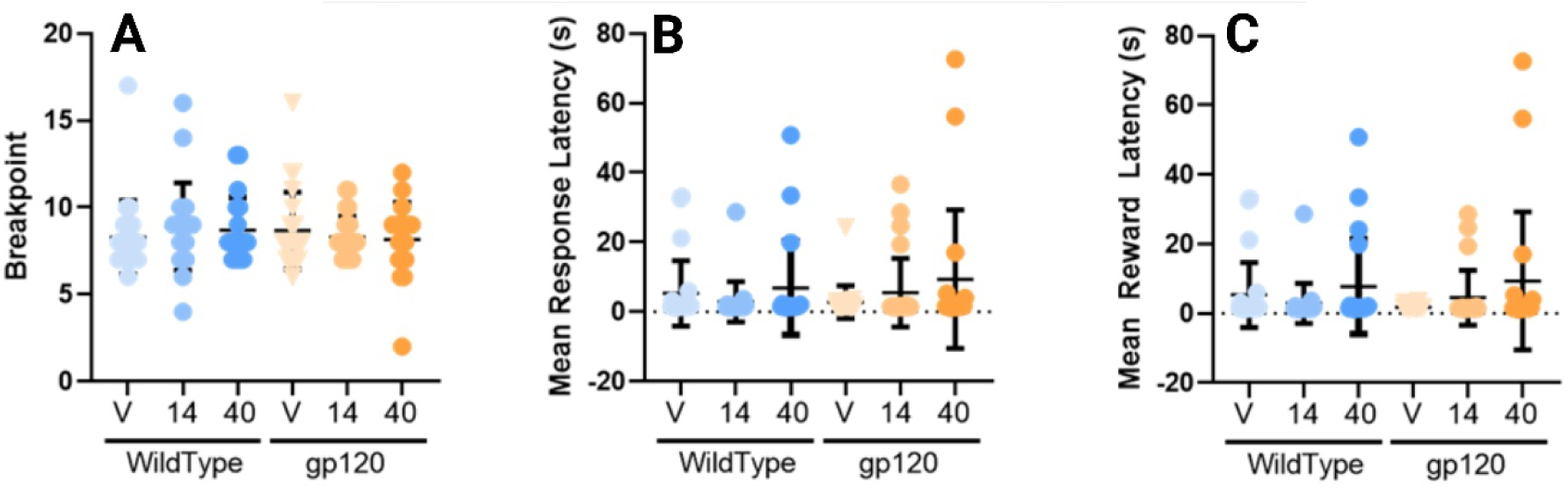
Task performance of groups in the PRBT task after chronic nicotine treatment. **A.** There were no effects of genotype or nicotine treatment on breakpoint. **B.** There were no effects of genotype or nicotine treatment on mean response latency. **C.** There were no effects of genotype or nicotine treatment on mean reward collection latency. Data presented as individual data-points, plus mean ± S.E.M.

#### Probabilistic Reversal Learning Task – Primary Measures

After testing in the PRBT, mice were tested in the PRLT after initial FR-1 retraining. <u>Switches:</u> We observed no significant main effect of sex, nicotine, or genotype (Fs<2.05, ns; Figure 3A). No sex*nicotine, sex*genotype, or sex*genotype*nicotine, interactions were observed (Fs<2.59, ns), but there was a trend toward a nicotine*genotype interaction [F(2,118)=2.75,*p*=0.068]. Given the *a priori* hypotheses, the impact of nicotine on each genotype was examined. Vehicle-treated WT and gp120-Tg mice did not differ in overall switches [F(1,42)=1.28,*p*=0.265]. No nicotine [F(2,60)=1.13,*p*=0.329], or nicotine*sex interaction [F(2,60)=1.05,*p*=0.355] was observed in WT mice. Despite increased switches in nicotine-treated gp120-Tg mice, there were no main effects of nicotine [F(2,58)=1.63, *p*=0.205], nor a sex*nicotine interaction [F(2,58)=0.548,*p*=0.581] in gp120-Tg mice. <u>%Target Responses:</u> A main effect of sex [F(1,118)=4.76,*p*=0.031] was observed, revealing males exhibited higher %Target Responses than females (Supplementary Figure 1G). No main effect of genotype or drug was observed (Fs<2.41, ns; Figure 3B). Importantly, a genotype*drug interaction was observed [F(2,118)=3.95,*p*=0.022]. *Post hoc* analyses revealed that nicotine did not affect WT mice [F(2,60)=0.92,*p*=0.403], but increased %Target Responses in gp120-Tg mice [F(2,58)=4.91,*p*=0.011], at both 14 and 40 mg/kg/day vs. vehicle (*p*=0.008 and 0.013, respectively; Figure 3B). This effect was observed despite no difference between vehicle-treated WT and gp120-Tg mice [F(1,42)=1.24,*p*=0.272].

**Figure 3.**
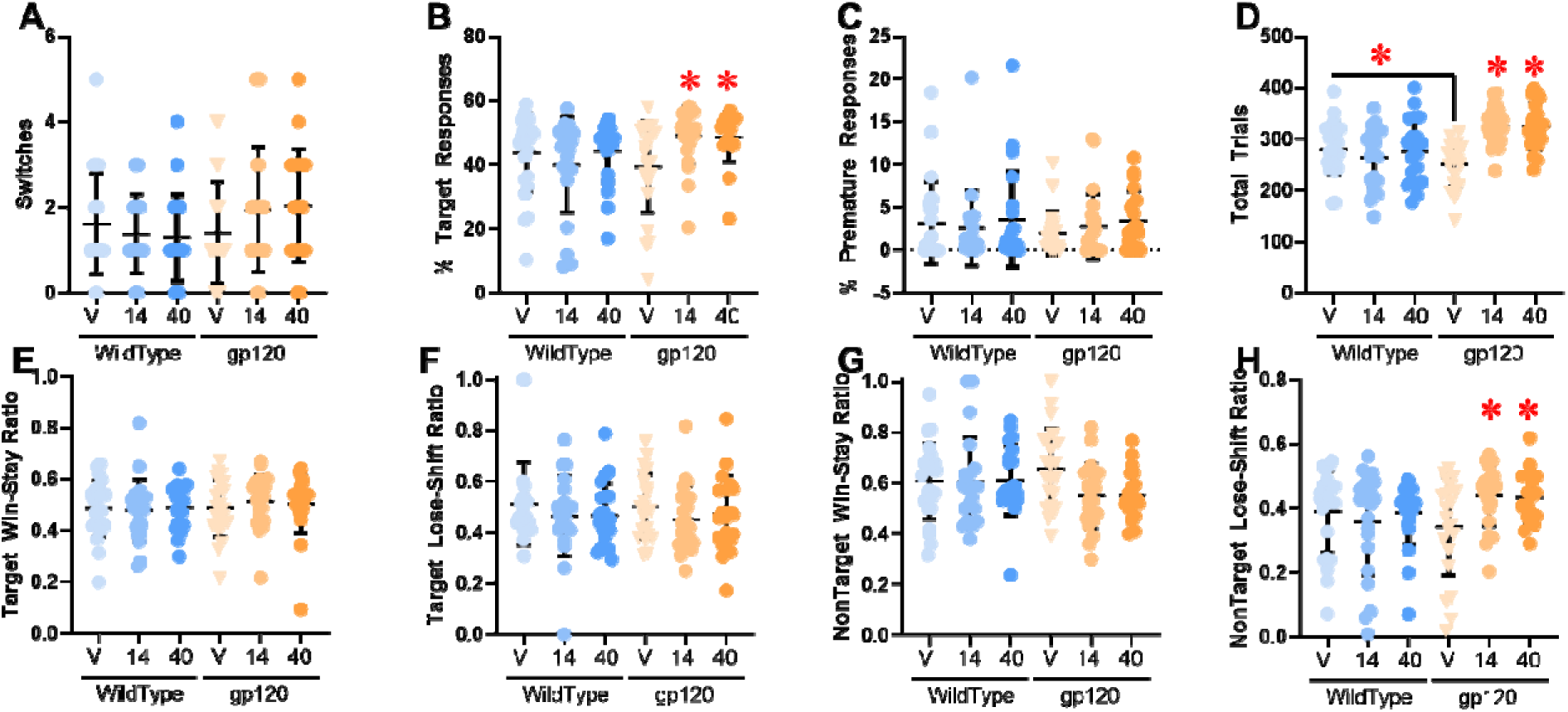
Task performance of groups in the PRLT during chronic nicotine treatment. **(A)** No group differences were observed in reversal learning as measured by reversal switches. **(B)** Chronic nicotine increased the percentage of target responses, selectively in gp120-Tg mice. **(C)** No group differences were observed in the percentage of premature responses. **(D)** Total trials were lower in gp120-Tg mice, and chronic nicotine increased total trials selectively in these mice **(E)**. No group differences were observed on Target Lose-Shift **(F)** or Non-Target Win-Stay **(G)** ratios. Chronic nicotine increased the Non-Target Lose-Shift Ratio, selectively in gp120-Tg mice **(H)**. Data presented as individual data-points, plus mean ± S.E.M. * denotes *p*<0.05 relative to vehicle (V)-treated mice within genotype unless explicit differences were made between genotypes.

#### Probabilistic Reversal Learning Task – Secondary Measures

<u>% Premature Responses:</u> A main effect of sex [F(1,118)=6.99,*p*=0.010] was observed, with female mice exhibiting fewer %premature responses than male mice. No main effect of genotype, assigned dose, nor interactions between sex*genotype, sex*assigned dose, genotype*assigned dose, or sex*genotype*assigned dose were observed (Fs<1.27, ns; Figure 3C). <u>Total Trials:</u> No main effect of sex, nor sex*genotype, sex*nicotine, or sex*genotype*nicotine, interactions were observed (Fs<1.22, ns). A main effect of genotype [F(1,118)=7.71,*p*=0.006], nicotine [F(2,118)=6.12,*p*=0.003], and a significant genotype*nicotine interaction [F(2,118)=10.76,*p*<0.001], were observed however (Figure 3D). *Post hoc* analyses revealed that vehicle-treated gp120-Tg mice completed fewer total trials than their WT littermates [F(1,42)=5.14,*p*=0.029]. As with other secondary measures, nicotine did not affect total trials of WT mice [F(2,60)=0.54,*p*=0.583], though it did increase total trials of gp120-Tg mice [F(2,58)=22.57,*p*<0.001], at both 14 and 40 mg/kg/day vs. vehicle (*ps*<0.001), perhaps unsurprising given that nicotine sped latencies to respond and collect reward (see below), in addition to improving %Target Responses resulting in fewer time-out periods.

#### Probabilistic Reversal Learning Task – Decision-Making Variables

<u>Target Win-Stay Ratio:</u> No main effect of sex, genotype, nicotine, nor sex*genotype, sex*nicotine, genotype*nicotine, or sex*genotype*nicotine, interactions were observed (Fs<0.52, ns; Figure 3E). <u>Target Lose-Shift Ratio:</u> No main effect of sex, genotype, nicotine, nor sex*genotype, sex*nicotine, genotype*nicotine, or sex*genotype*nicotine, interactions were observed (Fs<1.63, ns; Figure 3F). <u>NonTarget Win-Stay Ratio:</u> No main effect of sex, genotype, nicotine, nor sex*genotype, sex*nicotine, genotype*nicotine, or sex*genotype*nicotine, interactions were observed (Fs<2.26, ns; Figure 3G). <u>NonTarget Lose-Shift Ratio:</u> No main effect of sex, genotype, or nicotine, nor sex*genotype, sex*nicotine, or sex*genotype*nicotine, interactions were observed (Fs<2.52, ns). A genotype*nicotine [F(1,118)=3.15,*p*=0.046] interaction was observed however, with *post hoc* analyses revealing that while vehicle-treated WT and gp120-Tg mice did not differ [F(1,42)=1.26, *p*=0.269], and nicotine did not affect WT mice [F(2,60)=0.49,*p*=0.615], nicotine increased the lose-shift ratio of gp120-Tg mice when responding at the nontarget [F(2,58)=4.08,*p*=0.022], at both 14 and 40 mg/kg/day vs. vehicle (*p*<0.015 and *p*=0.026 respectively; Figure 3H).

#### Probabilistic Reversal Learning Task – Latency Measures

<u>Mean Target Latency:</u> No main effect of sex, genotype, nicotine, nor sex*genotype, sex*nicotine, or sex*genotype*nicotine, interactions were observed (Fs<2.3, ns). A genotype*nicotine interaction was observed however ([F(2,118)=4.42,*p*=0.014; Figure 4A). *Post hoc* analyses revealed that vehicle-treated gp120-Tg mice tended to be slower than their WT littermates [F(1,42)=3.04,*p*=0.089], and that while nicotine did not affect WT mice [F(2,60)=0.73,*p*=0.486], it sped target latencies of gp120-Tg mice [F(2,58)=10.13,*p*<0.001], at both 14 and 40 mg/kg/day vs. vehicle (*ps*<0.001s).

**Figure 4.**
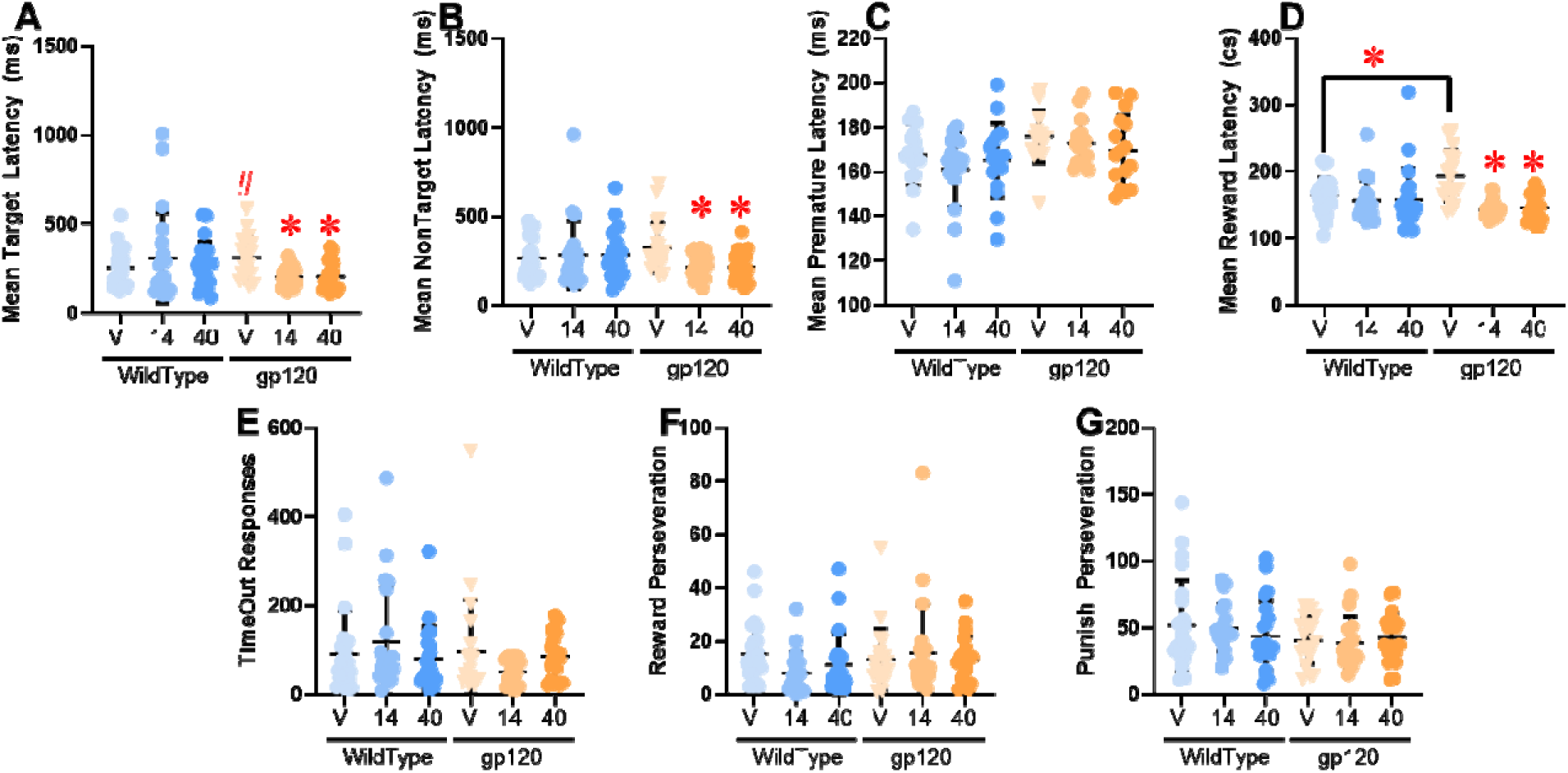
Chronic nicotine attenuated slowed latency deficits of gp120-Tg mice in the PRLT. **(A)** No group differences were observed in reversal learning as measured by reversal switches. **(B)** Chronic nicotine increased the percentage of target responses, selectively in gp120-Tg mice. **(C)** No group differences were observed in the percentage of premature responses. **(D)** Total trials were lower in gp120-Tg mice, and chronic nicotine increased total trials selectively in these mice **(E)**. No group differences were observed on Target Lose-Shift **(F)** or Non-Target Win-Stay **(G)** ratios. Chronic nicotine increased the Non-Target Lose-Shift Ratio, selectively in gp120-Tg mice **(H)**. Data presented as individual data points, mean ± standard error of the mean. * denotes *p*<0.05 relative to vehicle (V)-treated mice within genotype unless explicit differences were made between genotypes, # denotes *p*<0.1 relative to WT mice at V dose.

<u>Mean NonTarget Latency:</u> No main effect of sex genotype, nicotine, nor sex*genotype, or sex*genotype*nicotine, interactions were observed (Fs<1.7). A trend toward a sex*nicotine interaction [F(2,118)=2.56,*p*=0.081], and a significant genotype*nicotine interaction [F(2,118)=3.66,*p*=0.029], were observed however (Figure 4B). *Post hoc* analyses revealed that vehicle-treated gp120-Tg mice did not differ from their WT littermates [F(1,42)=2.48,*p*=0.123]. As with mean target latency, nicotine did not affect nontarget latency of WT mice [F(2,60)=0.83,*p*=0.452], but it sped nontarget latencies of gp120-Tg mice [F(2,58)=7.76,*p*=0.001], at both 14 and 40 mg/kg/day vs. vehicle (*p*=0.002 and *p*=0.003 respectively).

<u>Mean Premature Latency:</u> No main effect of sex [F(1,118)=0.30,*p*=0.585], nicotine [F(2,118)=1.17,*p*=0.315], nor sex*nicotine [F(2,118)=2.09,*p*=0.130], genotype*nicotine [F(2,118)=0.81,*p*=0.448], or sex*genotype*nicotine [F(2,118)=0.19,*p=*0.830], interactions were observed. A main effect of genotype [F(1,118)=6.67, *p*=0.011], and a sex*genotype interaction [F(1,118)=4.04, *p*=0.048], was observed, with WT mice exhibiting faster premature latencies than the gp120-Tg mice (Figure 4C). *Post hoc* analyses revealed this effect was primarily in female mice, although no effect of genotype in either sex was observed (*p*s>0.1).

<u>Mean Reward Latency:</u> No main effect of sex, genotype, nor sex*genotype, sex*nicotine interaction, or sex*genotype*nicotine, interactions were observed (Fs<1.33, ns). A main effect of nicotine [F(2,118)=11.75,*p*<0.0001], and a significant genotype*nicotine interaction [F(2,118)=6.79,*p*=0.002], were observed however (Figure 4D). *Post hoc* analyses revealed that vehicle-treated gp120-Tg mice were slower to collect rewards than their WT littermates [F(1,42)=9.16,*p*=0.004]. As with mean target and nontarget latencies, nicotine did not affect reward latency of WT mice [F(2,60)=0.28,*p*=0.761], it did speed target latencies of gp120-Tg mice [F(2,58)=25.80,*p*<0.001], at both 14 and 40 mg/kg/day vs. vehicle (*ps*<0.001).

#### Probabilistic Reversal Learning Task – Secondary Outcomes

<u>Time Out Responses:</u> No main effect of sex, genotype, nicotine, nor sex*genotype, sex*nicotine, genotype*nicotine, or sex*genotype*nicotine, interactions were observed (Fs<2.25, ns; Figure 4E). <u>Reward Perseverative Responses:</u> A main effect of sex [F(1,118)=9.89,*p*=0.002], with female mice exhibiting fewer perseverative responses after gaining a reward than male mice. No main effect of genotype, nicotine, nor sex*genotype, sex*nicotine, genotype*nicotine, or sex*genotype*nicotine, interactions were observed (Fs<1.79, ns; Figure 4F). <u>Punish Perseverative Responses:</u> A main effect of sex [F(1,118)=8.38,*p*=0.005] was observed, with female mice exhibiting fewer perseverative responses after gaining being punished than male mice. A trend effect of genotype [F(1,118)=2.95,*p*=0.088] was also observed, which revealed slightly higher perseverative responses in WT mice. No effect of nicotine, nor sex*genotype, sex*nicotine, genotype*nicotine, or sex*genotype*nicotine, interactions were observed (Fs<1.4, ns; Figure 4G).

#### Iowa Gambling Task

To confirm validity of the task, we evaluated performance across three trial blocks on the IGT and observed a significant main effect of block on difference score [F(2,348)=397.0,*p*<0.001], in addition to a block*genotype*drug interaction [F(4,348)=3.1,*p*<0.05]. *Post hoc* analyses did not reveal specific within group effects however. No effect of block by drug (F<1.2, ns), or block by genotype (F<1, ns) interaction was observed, therefore data were collapsed across blocks for further analysis. When data were collapsed across blocks, we observed a trend effect of sex [F(1,118)=2.9,*p*=0.097] which revealed that females exhibited less risk preference than male mice.

With the data collapsed across blocks we observed a significant genotype*drug interaction for difference score ([F(1,118)=4.6,*p*=0.015]; Figure 5A). There was also a trending difference score genotype*drug interaction for punishment duration (F(1,118)=2.8,*p*=0.070; Figure 5B). No gene or drug effects, nor their interactions were observed on any other measure (Fs<2.3, *p*s>0.1, see Supplemental Table 1). *Post hoc* analyses revealed no effect of drug in WT mice (F<1.7, ns). In contrast, nicotine tended to increase difference score (F(2,58)=2.6,*p*=0.09) and reduce punishment duration [F(2,65)=2.6,*p*=0.098] in gp120-Tg mice, with no effect in any other measure (F<1.6, ns; Figures 5C-H). Further analyses revealed that gp120-Tg mice treated with 40 mg/kg/day exhibited higher difference scores than those treated with vehicle (*p=*0.071), with no other effects or trends observed. One sample t-test revealed that in-terms of difference score, WT mice treated with vehicle exhibited a difference score significantly higher than chance (t=1.948,*p*<0.05), as did gp120-Tg mice treated with 40 mg/kg/day (t=2.4,*p*<0.05), with no other group differing from chance (0).

**Figure 5.**
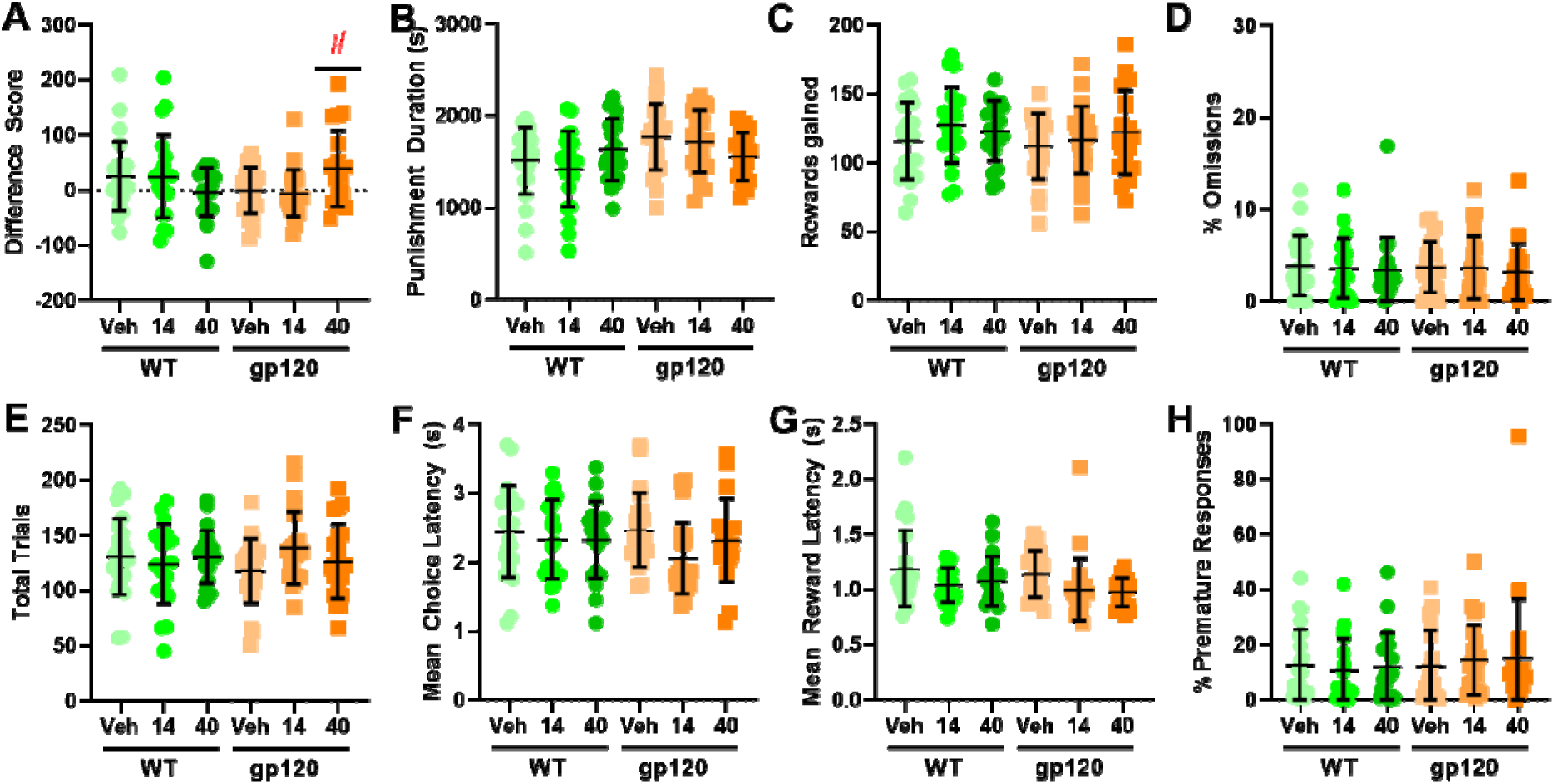
Nicotine optimized IGT performance in gp120-Tg but not WT mice. **(A)** High dose nicotine (40 mg/kg/day) tended to increase difference score in gp120-Tg mice only. **(B)** There was trend towards a genotype*drug interaction for punishment duration, with a reduction observed in gp120-Tg mice. **(C)** There were no effects of genotype or nicotine treatment on number of rewards gained, **(D)** % omissions, **(E)** total number of trials, **(F)** mean choice latency, **(G)** reward collection latency, or **(H)** %premature responses. Data represents individual data, plus mean ± standard error of the mean. # = p<0.1 vs V within gp120-Tg mice.

#### Nicotine reduced Iba1 staining in both gp120-Tg and WT littermate mice

After final testing at day 28, brains were immediately removed for analysis for microglial activation (Iba1 staining). A trend toward a drug effect was observed [F(2,21)=3.3,*p*=0.056], revealing that nicotine at the highest dose reduced Iba1 levels irrespective of genotype (Figure 6).

**Figure 6.**
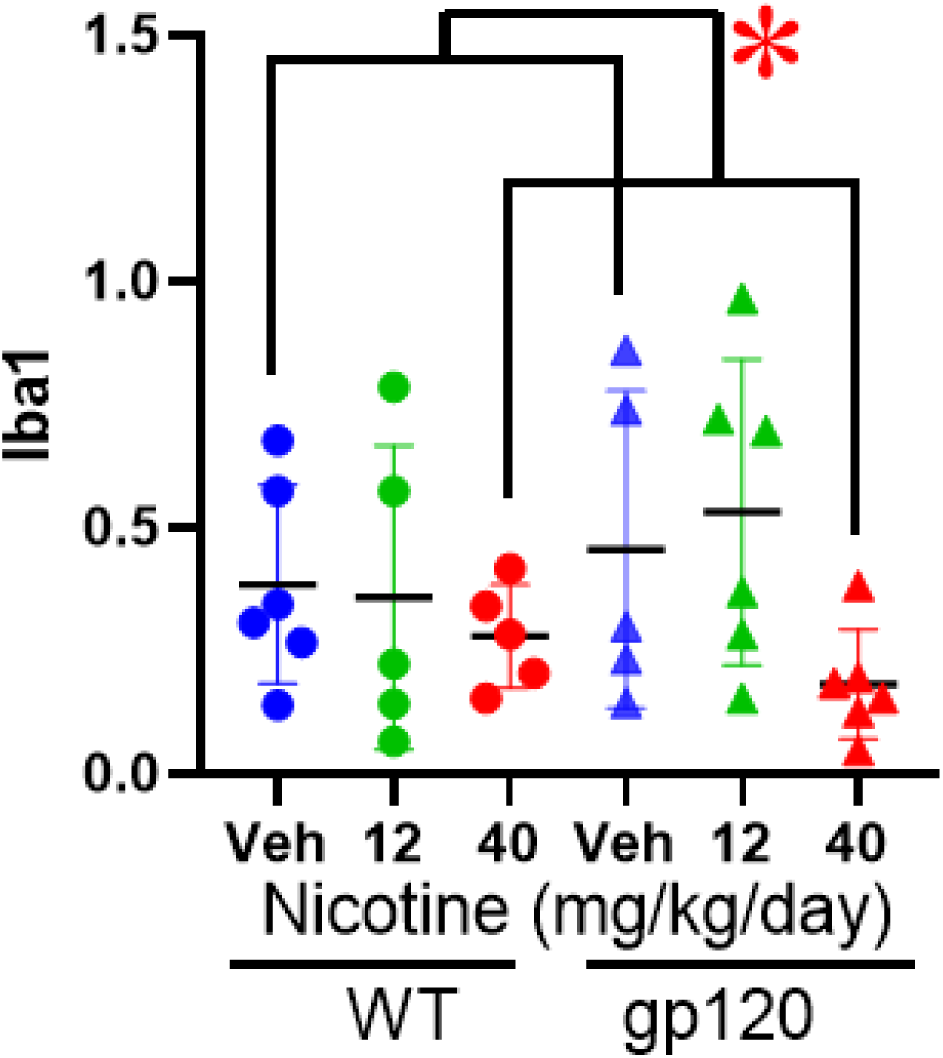
Chronic nicotine lowered neuroinflammation as measured by Iba1 levels. Chronic nicotine lowered Iba1 levels in mice irrespective of genotype. Data represents individual data points, plus mean ± SEM. * =*p*<0.05

## Discussion

Using a battery of translationally relevant cognitive tasks, we provide a characterization of the motivational and decision-making profile in the gp120-Tg mouse model of HIV pathology. While gp120-Tg mice exhibited similar PRBT and IGT performance compared to WT controls at baseline, they exhibited altered PRLT performance, consistent with impaired learning in PWH. Importantly, chronic nicotine improved PRLT and IGT performance in gp120-Tg mice, not in their WT littermates. Hence, nicotine induced a shift towards a “safer” profile of decision-making under probabilistic and risky conditions. Finally, we demonstrated that chronic nicotine treatment reduced neuroinflammation in gp120-Tg and WT mice, possibly providing a mechanism for improved cognition and insight into the therapeutic benefits of nicotine use in PWH.

### Associative learning and motivation in gp120-Tg mice

We first trained gp120-Tg mice on fixed ratio operant responding and the PRBT. Similar to previous reports, gp120-Tg mice exhibited no deficits in acquiring fixed ratio responding or breakpoint, suggesting intact associative learning and motivational processes [58, 59]. These findings are consistent with a lack of PRBT deficits in the HIV1-tg rat model [60] (though see [61]). PWH exhibit impaired learning [62–64], and gp120-Tg mice have previously showed deficits during the initial acquisition phase of the attention set-shifting task (ASST) [65]. The acquisition of the ASST requires making associations using multiple domains of sensory information (i.e., odor, texture), suggesting that more challenging paradigms are likely necessary to observe learning impairments in gp120-Tg mice. Previous studies indicate gp120-Tg mice exhibit increased sensitivity to rewards, including for saccharin solution and methamphetamine, as measured by increased consumption and stronger conditioned place preference, respectively [59]. Using a highly palatable strawberry milkshake reward in our paradigms, we did not observe any indication of increased reward drive, neither on the PRBT nor in ancillary measures such as response or reward collection latency. Hence, heightened reward sensitivity did not impact their effortful motivation to work for a reward. Finally, as observed previously in control mice [66], chronic nicotine also did not affect breakpoint for food rewards in either genotype, suggesting motivation to work for palatable rewards was unaffected by genotype or drug.

### Reinforcement learning and cognitive flexibility

We next assessed gp120-Tg mice on the PRLT, providing an evaluation of reinforcement learning and cognitive flexibility. PWH exhibit impaired learning in traditional paradigms such as the Wisconsin card sorting task albeit the PRLT has yet to be extensively tested in PWH. HIV1-tg rats exhibit normal, or even improved initial probabilistic learning but normal levels of reversal learning in a within-session paradigm [67, 68] while they exhibit impaired reversal learning when tested across sessions using deterministic instead of probabilistic conditions [69, 70]. Interestingly, gp120-Tg mice did not exhibit impaired reversal learning nor abnormalities in responses to probabilistic feedback, aligning with the HIV1-tg rats. Gp120-Tg mice demonstrated a significant reduction in the overall trials completed and a trend towards slowed correct response latency however, potentially revealing a speed-accuracy trade-off indicative of impaired performance. Slowed response latencies and reduced trial counts were also repeatedly observed in HIV1-tg rats which may also reflect a speed-accuracy tradeoff [67, 68]. Hence, potential sub-optimal choices in HIV model animals could be attenuated by taking longer to choose at the cost of completing more trials [60, 71]. Such slowed reaction times are seen in PWH performing computerized tasks [72] likely reflecting speed-accuracy trade-offs. Given that HIV1-tg rats overexpress gp120-Tg as do the gp120-Tg mice, this protein may drive impaired complex cognitive performance.

While the impact of gp120-Tg on learning and cognition may have been modest, we observed numerous interactions between genotype and chronic nicotine treatment, with nicotine effects specific to gp120-Tg mice. Both nicotine doses attenuated the baseline deficit in %target responses and sped response latencies to target and non-target stimuli. Nicotine-induced improvement in % target responses likely arise from their increase in non-target lose-shift – mice were more likely shift after being punished on the non-target options. These findings suggest nicotine improved reinforcement learning in gp120-Tg mice by heightening sensitivity to losses. Furthermore, nicotine attenuated the processing speed deficits in these mice. Acute nicotine treatment previously improved reinforcement learning in PRLT performance in WT mice [73], with evidence confirming that nicotine improved attention and reaction times in rodents and humans [7, 74–78]. While the effects of nicotine consumption in PWH produce conflicting results in regard to some cognitive functions [20, 79], these data indicate nicotine was effective at alleviating deficits in learning and/or processing speed in gp120-Tg mice. Lastly, although motivation appeared intact on the PRBT, gp120-Tg mice exhibited a baseline slowed reward collection latency compared to WTs on the PRLT, typically indicative of reduced reward motivation. Although this effect appeared to be transient, it was also alleviated with nicotine treatment at both doses. Hence, nicotine may more specifically attenuate reinforcement learning deficits in gp120-Tg mice.

### Risk-based decision-making in gp120 mice

The IGT enables decision-making to be evaluated under conditions of response-outcome uncertainty (risk). Gp120-Tg mice exhibited overall worse performance as measured by difference score in the IGT as seen in PWH [80, 81] albeit non-significantly. Importantly, nicotine increased difference score specifically in gp120-Tg mice, reflecting a shift towards a “safer” response strategy [54, 82]. Hence, nicotine use in PWH drives better decision making under risk in addition to better reinforcement learning. These findings have implications for the therapeutic potential of nicotine for targeting poorly optimized decision-making in PWH.

Previously, chronic nicotine treatment (0.25 μL/h over 28 days) increased the number of successful trials and reduced choice latency in a deterministic task [83]. In a probabilistic paradigm nicotine shifted responding to the most consistently rewarded option [83] consistent with our observations of increased target responding, lose-shift behaviors, and increased difference scores. Thus, our findings are in agreement that chronic nicotine treatment facilitated adaptive decision making in accordance with the probability of reward.

### Reduced neuroinflammation potentially underlying nicotine-mediated effects in gp120-Tg mice

Nicotine selectively improved task performance in gp120-Tg mice even in the absence of baseline differences between genotypes. To determine one potential mechanism, we tested whether nicotine differentially altered physiological measures between WT and gp120-Tg mice by evaluating Iba-1 levels following cognitive testing. Iba-1 is an indicator of microglia activation, thus providing an assessment of differences in neuroinflammation [84]. PET imaging studies have reported correlations between increased markers of microglia activation and poorer cognitive performance in PWH [85, 86]. Moreover, we recently demonstrated that PWH that recently smoked had improved attention and reduced indicators of neuroinflammation compared to non-smoking PWH [15]. Hence, addressing neuroinflammation could be key to improving cognition in mouse models and PWH.

Interestingly, we did not observe baseline differences in Iba-1 levels between genotypes. While this finding is in agreement with a previous assessment of Iba-1 levels in gp120-Tg mice [87], and in PET imaging studies at baseline [39], increased Iba-1 levels have been reported in transgenic gp120-Tg mice [88] and in the initial days following treatment with gp120-Tg [89]. This difference may have been a result of the low sample sizes in the current studies. Increased microglial activation (measured by the translocator protein) was observed in *in vivo* PET imaging studies in gp120-Tg mice relative to their WT littermates only in response to the lipopolysaccharide (LPS) immune challenges, indicating this model recapitulates abnormalities in neuroinflammatory responses present in PWH [39]. We observed however, that chronic nicotine treatment reduced Iba-1 levels in both WT and gp120-Tg mice. Given that our findings that both the cognitive and physiological benefits of chronic nicotine were selective to the gp120-Tg mice, it is possible that reduced neuroinflammation was associated with improved task performance. Despite nicotine-induced reduction in Iba1 staining in WT mice, improved cognition was not seen in these mice, potentially due to performance already at optimal levels in these tasks. Future experiments should expand upon the method and/or target for measuring the neuroinflammatory response in gp120-Tg mice, which could provide further insight for the mechanism underlying the cognitive enhancing effects of nicotine in this model of HIV-NCI.

### Summary

In conclusion, gp120-Tg mice exhibited intact motivation and modest deficits in reinforcement learning and risk-based decision making. Importantly, these deficits were attenuated with chronic nicotine treatment at both doses. In gp120-Tg mice, nicotine also altered the response strategies on the PRLT and IGT, shifting mice towards a bias for the most consistently rewarded option and reducing risky decision-making. The specificity of cognition enhancing effects of nicotine in gp120-Tg mice suggest it is acting on underlying mechanisms that are operating differentially between gp120-Tg and WT mice. Although we observed no group differences in the Iba-1 marker of microglia activation under saline, nicotine treatment significantly reduced Iba-1 levels in both groups, suggesting that reductions in neuroinflammation may be associated with the cognitive effects observed in gp120-Tg mice. Together, these data provide a translationally relevant assessment of the effects of nicotine in the context of HIV-relevant pathology, suggesting nicotine may confer therapeutic potential as a compound for targeting risky decision-making and underling neuroinflammation.

