## Supplementary files for "Chronic nicotine treatment enhances cognition and reduces neuroinflammation in the gp120 transgenic mouse model of neuroHIV"

**SUPPLEMENTARY MATERIAL**

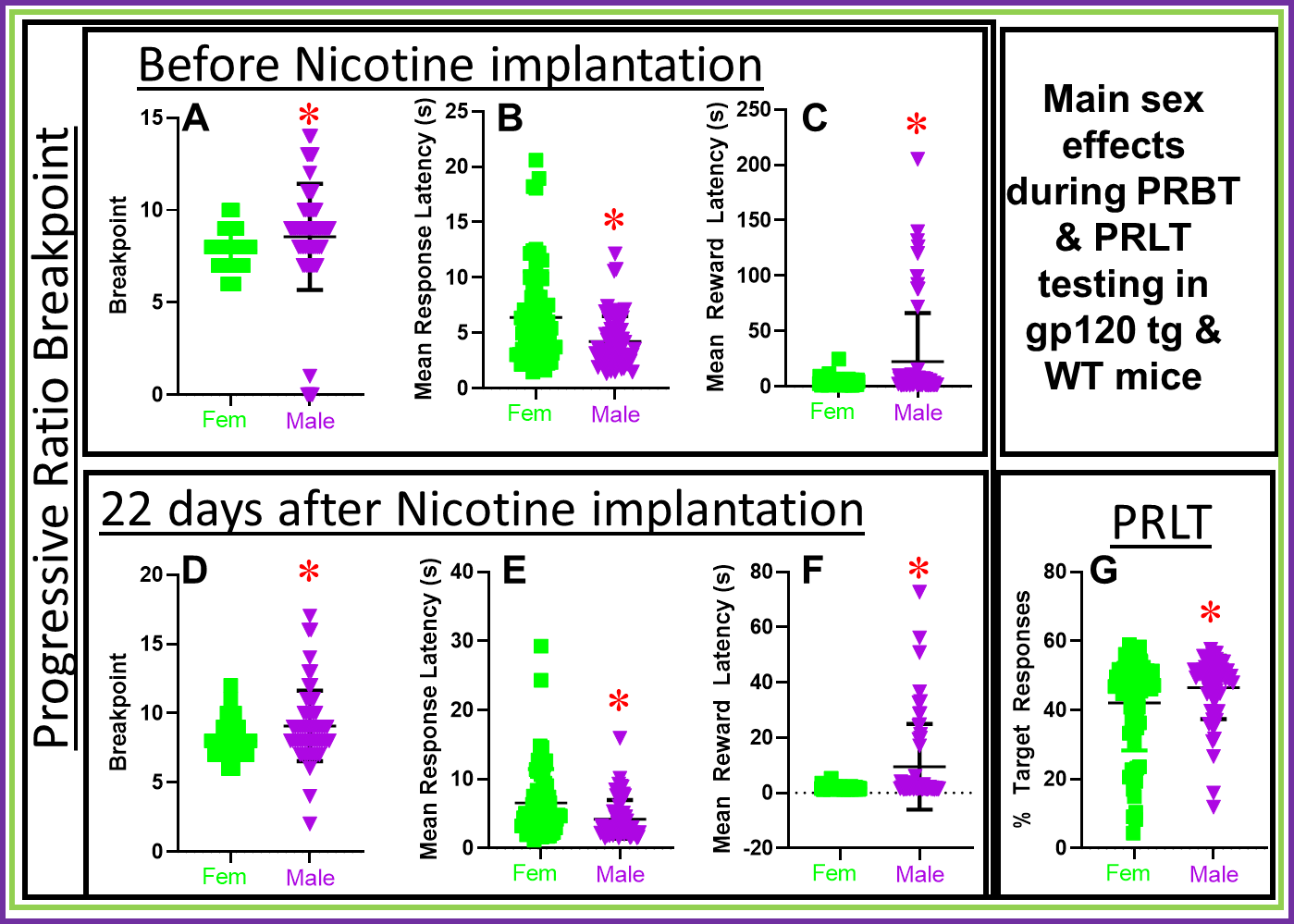

**Supplemental Figure 1: Sex differences across tasks throughout testing.** Prior to nicotine implantation male mice exhibited elevated breakpoint (**A**) alongside faster response latency (**B**). and slowed latency to collect rewards (**C**) relative to female mice. These differences were consistent even testing more than three-weeks later (**D-F)**. Males also exhibited elevated %target responses during PRLT performance (**G**).

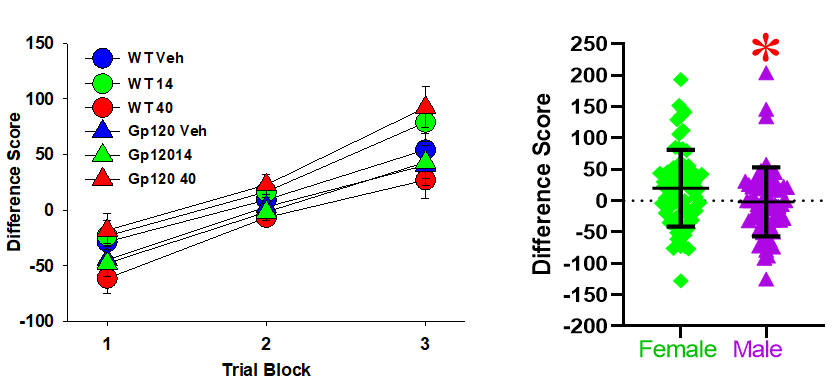

**Supplemental Figure 2. Within-session learning of the IGT of gp120-Tg and wildtype (WT) littermate mice, with males choosing risker rewards than females.** Both gp120-Tg and WT littermate mice exhibited a shift toward preferring the safe choices over time (increased difference score) from below to above chance (0) levels (left), while male mice exhibited poorer IGT performance relative to females (right). ***** denotes *p*<0.05 relative to females.

**Supplemental Table 1.** Impact of nicotine treatment on secondary measures in the Iowa Gambling Task Performance. Denotes F values followed by p value.

| Factor | DoF | Difference Score | Rewards | Punish Duration | %Omissions | MCL | MRL | %Prems |
| --- | --- | --- | --- | --- | --- | --- | --- | --- |
| Gene | 1,118 | .6, .45 | .002, .97 | .002, .97 | .41, .53 | .27, .61 | .03, .86 | .002, 0.96 |
| Drug | 2,118 | .15, .87 | .76, .47 | .11, .89 | .81, .45 | 1.1,.35 | 2.3, .11 | .25, .78 |
| Sex | 1,118 | ***2.9, .097*** | 2.5, .123 | .048, .83 | .13, .73 | .026, .87 | 2.5, .12 | .009, .93 |
| G*D | 1,118 | **4.6, .015** | 1.8, .17 | ***2.8, .070*** | .20, .819 | 1.4, .25 | .52, .60 | .08, .92 |
| G*S | 1,118 | 1.2, .29 | .01,.91 | .02, .88 | .02, .89 | .88, .35 | .02, .89 | .47, .50 |
| S*D | 1,118 | 1.3, .28 | 1.4, .25 | 2.0, .14 | 1.8, .17 | .44, .64 | 1.4, .26 | 1.1, .34 |
| G*S*D | 2,118 | .78, .47 | 1.0, .38 | 1.4, .25 | .16, .86 | 1.2, .32 | .94, .40 | .97, .39 |
